# The Effect of Arm Restriction on Dynamic Stability and Upper Body Responses to Lateral Loss of Balance During Walking: An Observational Study

**DOI:** 10.1101/2023.09.11.557158

**Authors:** Uri Rosenblum, Adi Lavi, Arielle Fischer, Yisrael Parmet, Amir Haim, Shirley Handelzalts

**Author notes:** Equal contribution (Amir Haim, Shirley Handelzalts). Corresponding author: Shirley Handelzalts PhD PT, Department of Physical Therapy, Recanati School for Community Health Professions, Faculty of Health Sciences. Ben-Gurion University of the Negev, Israel. **Declaration of conflicting interests:** The authors declare no conflict of interest.

## Abstract

When losing balance, upper-body movements serve as mechanical aids to regain stability. However, it remains unclear how these movements contribute to dynamic stability during recovery from a lateral loss of balance while walking with arm restriction. We aimed to 1) quantify the effect of arm restriction on gait stability and upper-body velocities, and 2) characterize upper-body kinematic strategies in response to lateral surface translations under different arm restriction conditions. Healthy adults were exposed to lateral surface translations while walking on a computerized treadmill under three conditions: ‘free arms’, ‘1-arm restricted’ and, ‘2-arms restricted’. Dynamic stability and upper-body velocities for the first step after perturbation onset were extracted. We found decreased dynamic stability in the sagittal plane and increased trunk velocity in the ‘2-arm restricted’ condition compared to the ‘free arms’ condition. Head and trunk movements in the mediolateral plane were in opposite directions in 44.31% of responses. Additionally, significant trunk velocities were observed in the opposite direction to the perturbation-induced loss of balance. Our results support the contribution of increased upper-body velocities to balance responses following arm-restricted walking perturbations and suggest that the ‘2-arm restricted’ condition may be utilized as a perturbation-based balance training, focusing on head and trunk responses.

## Background

Reactive balance responses are critical for maintaining stability following a sudden loss of balance. While extensive research has delved into understanding lower limb reactions to balance disruptions (1–5), the contribution of upper body responses in restoring balance are not fully understood. Increasing the understanding of upper body responses to balance perturbations is vital for identifying rehabilitation training targets for fall prevention.

To avoid falls, regulating the velocity of the upper body (i.e., arms, head, and trunk) is crucial due to their combined mass which constitutes about two-thirds of the total body mass (6), their height above the feet (i.e., base of support, BoS), and their impact on the movement of the center of mass (CoM). Additionally, stabilizing the head on the trunk is essential for balance control (7–9).

During unperturbed walking, trunk and arm motion has been shown to enhance stability by counteracting the lower body’s angular momentum (10–13). Conversely, studies have shown that walking with arms held may improve stability by increasing trunk inertia, which limits center of mass (CoM) displacement (14,15). When introducing perturbations during walking, the lack of arm motion (i.e., restriction) has affected gait parameters (14,16) and led to a slower return towards the normal gait pattern compared to walking with normal arm motion (14). In response to perturbations upper extremity strategies have been characterized as a protective strategy against injury, such as positioning the upper extremities to absorb impact energy with the ground or as a recovery strategy to stabilize the body by either reach-to-grasp or assist in counter balancing and decelerating the falling center of mass (15,17,18). A recent systematic review (19) highlighted differences in upper extremity responses to different perturbation types (e.g., unexpected perturbations during standing and slips and trips during walking) between young and older adults. For example, directional differences in upper extremity responses were demonstrated: older adults tended to move their arms in the same direction as perturbation while young adults tended to move their arms opposite to the direction of perturbation. These directional differences suggest an attempt to arrest the fall at impact in older adults (i.e., to brace against impact) and an attempt to restore an upright position by decreasing fall directed CoM displacement in young adults (i.e., to restore balance and prevent a fall) (20,21). Other studies focusing on slips reported bilateral arm flexion to counter the backwards loss of balance (18) and to reduce trunk extension velocity, which aids in regaining balance (22). While most studies have focused on slips that induce a backward loss of balance (18,22–24) or trips (20,25) that induce a forward loss of balance, in the real world losses of balance can occur in all directions. In fact, the literature shows that mediolateral (ML) loss of balance is more challenging to older adults compared to anterior-posterior loss of balance. It is frequently associated with more stepping errors (e.g., leg collisions) (26,27), with altered first step characteristics and postural movements of the trunk (27) and greater arm reactions in older adults (26,27). Finally, balance reactions to ML loss of balance were found to be predictors of falls (28).

Margins of stability (MoS), i.e., the distance between the extrapolated CoM (*XcoM*, see eq.1 in Methods) and the border of the BoS (29), is a measure of dynamic stability during both unperturbed and perturbed walking conditions (30,31). Previous research has demonstrated that MoS is a key determinant in identifying balance outcomes (i.e., loss of balance) following perturbations (1,32,33). Moreover, significant differences in MoS during reactive steps to perturbations have been observed between older adults with and without a history of falls (34).

To date, limited data exist regarding upper body responses to lateral perturbations during walking. Additionally, the potential effects of restricting the upper extremity movements on dynamic stability (i.e., MoS) and upper body reactive strategies in response to lateral perturbations have not been experimentally tested. This is particularly important given that humans are most dynamically unstable in the frontal plane (35–37) and that daily activities (e.g., talking/texting messages on the phone or carrying a bag while walking) or certain orthopedic/neurological conditions (e.g., upper limb fractures, stroke) may impose limitations on upper extremity responses. A previous study that employed an arm restriction approach to investigate upper body responses to slips in young adults demonstrated a significantly higher rate of falls when both arms were restricted (23). It was suggested that the increased fall frequency in the arm restricted condition resulted from the inability to initiate an effective alternative strategy to control the CoM excursion and recover balance. This alternative strategy should be utilized by other body segments (e.g., trunk and head). Since walking introduces inertia in the AP direction while ML surface translation introduces acceleration in the ML direction, we should find mixed directions of acceleration for the different body segments. However, this study did not assess body kinematics and upper body segments’ movement strategies used to recover balance (e.g., whether the body segments’ velocity was in the same direction as the perturbation-induced fall or in the opposite direction), limiting our understanding of the underlaying mechanisms. Therefore, the objectives of the current study were twofold: 1) to quantify the effect of arm restriction on dynamic stability and upper body kinematics in response to lateral surface translations during gait, and 2) to characterize upper body kinematics and strategies in response to lateral surface translations under different arm-restriction conditions. We hypothesized that 1) greater arm restriction will lead to decreased dynamic stability and increased upper body segment velocities to compensate for restricted arm movement, and 2) head and trunk velocities will be greater in the mediolateral direction to counteract the initial velocity in the direction of balance loss (i.e., the perturbation induced velocity).

## Methods

### Participants

Fourteen healthy young adults (7 females; aged 35±2.3 years), without any neurological, vestibular, or orthopedic conditions that may affect their balance and/or gait participated in the study. All participants signed an informed consent form prior to participating in the study. The study was approved by the institution review board at Loewenstein Rehabilitation Medical Center (LRMC) (approval number LOE-14-0021).

### Procedure

Assessments were conducted at the Gait Recovery Laboratory in LRMC. Participants walked on a computerized treadmill system with a horizontal movable platform (Balance Tutor, MediTouch LTD, Tnuvot, Israel) at their preferred walking speed, determined from the average of two overground 10-meter walk tests. Participants walked under three conditions, each lasting 120 seconds, in the following order: 1) unrestricted upper body movement, ‘free arms’ condition; 2) the dominant arm, determined by asking the participant which hand they use for writing, restricted with a sling, ‘1-arm restricted’ condition; 3) both arms restricted with slings, ‘2-arm restricted’ condition (Figure 1). In each condition, participants were exposed to four lateral surface translations (rightward, leftward, leftward, and rightward) with a platform displacement of 12 cm, a velocity of 42 cm/sec and an acceleration of 145 cm/sec^2^). They were not informed about the timing or direction of the perturbations. Perturbations were introduced randomly every 30-45 seconds to reduce predictability. Participants were instructed to react naturally to prevent falling during the surface translations. A safety harness attached to an overhead support protected the participants in case of a fall, but it did not support any body weight nor restricted their movements during walking.

**Figure 1.**
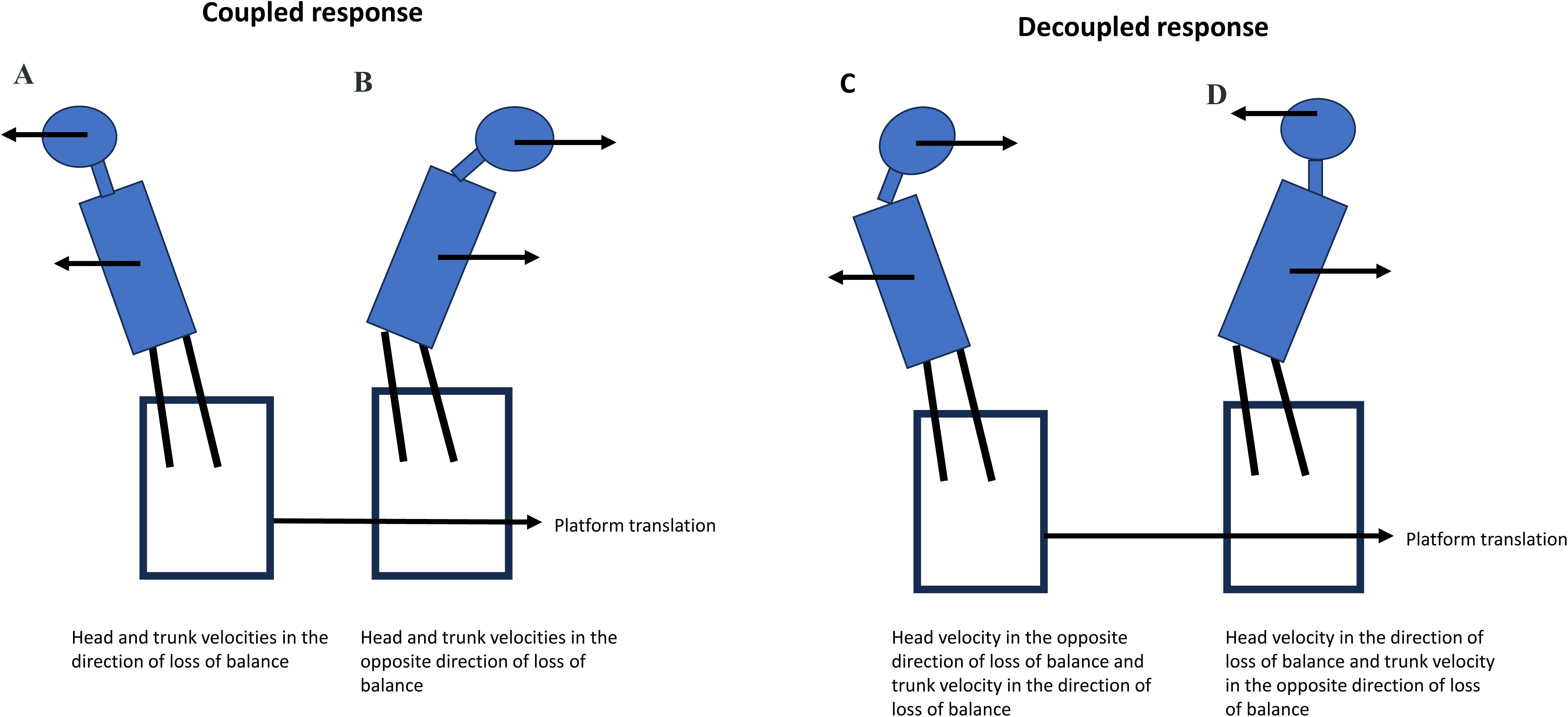
Experimental setup.

#### Kinematic data collection and analyses

Thirty-four reflective markers were strategically placed on anatomical landmarks in accordance with the Plug-In-Gait marker set (full body Plug in Gait model, Vicon Motion Systems, Oxford, UK). Three additional markers were affixed to the platform to synchronize platform movement data with kinematic data. Eight Vicon infrared motion capture cameras (Vicon Motion System, Oxford, UK) recorded kinematic data at a sampling rate of 120 Hz. Prior to analysis, motion data were preprocessed and analyzed using Vicon Nexus software (Version 2.0) and checked for marker trajectory disruptions. Then, a 3D model was generated using the software built-in function and the body CoM was extracted. The marker data were low-pass filtered using a zero-lag, fourth-order Butterworth filter with a cut-off frequency of 8 Hz. Gait events and parameters were computed using custom MATLAB scripts (version R2022b, MathWorks, Natick USA). Preprocessed data were used to identify the initial contact, defined as the local maxima of the heel marker position in the anterior-posterior (AP) direction (38,39). Gait spatiotemporal parameters (e.g., step length and width) were calculated from the marker data (see mean values in Table S1 in *supplementary materials*).

### Kinematic outcome measures

All outcome measures were calculated for the first step after perturbation onset, as the main balance responses, particularly those involving the upper limbs, are most prominent in the early recovery phase following perturbations (20,40).

### Margin of stability in the mediolateral (ML) and anterior posterior (AP) directions

To assess dynamic stability, the Margin of stability (MoS) in the anterior-posterior (AP) and mediolateral (ML) directions were calculated using equations 1 and 2, based on (29):

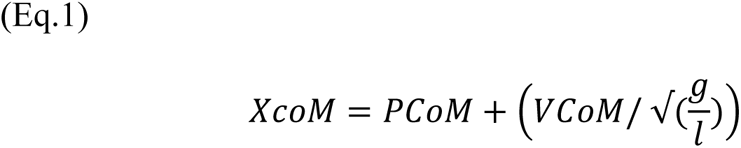

Where *XcoM* is the extrapolated center of mass, P_CoM_ is the AP or ML component of the projection of the center of mass on the ground. V_CoM_ is the CoM-velocity in the respective direction subtracted by the perturbation platform velocity. The term √(g/l) presents the eigen frequency of an inverted pendulum system with a leg of length l.

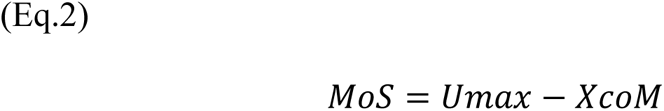

Where MoS_x_ specifies the MoS in the AP or ML direction (from herein the term **MoS_AP** and **MoS_ML** will be used), Umax is the anterior or lateral boundary of the base of support (BoS) defined as the toe marker in AP direction and lateral malleolus in the ML direction, and the XcoM is the position of the XcoM in the relevant direction.

### Area under the curve (AUC) of head and trunk velocities

To compute the velocity of the head and trunk we first determined their positions. The head position was determined as the average of the four head markers in 3D space. Trunk position was determined as the average position of the sternum, C7 and T10 markers in 3D space. The first derivative of the preprocessed marker position of the trunk and the head were calculated to obtain velocity using equations 3 and 4:

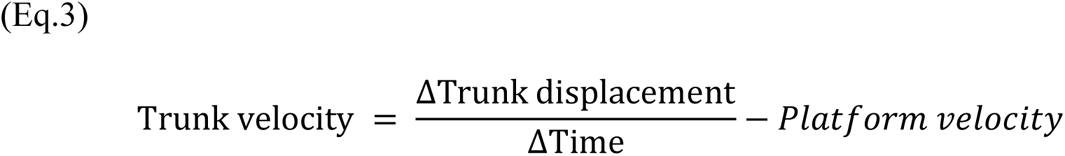

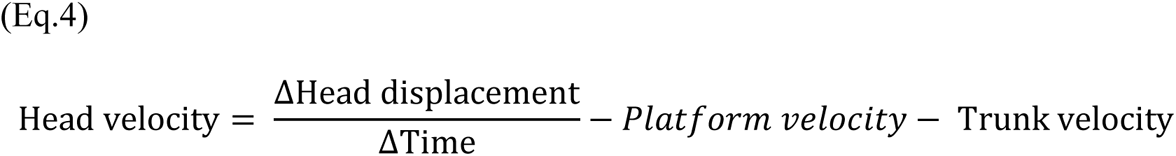

Where platform velocity is the velocity of the platform during the perturbation.

Head and trunk velocity vectors were interpolated to 101 samples to normalize step time after perturbation to 100%. We used the *absolute* and *true* values of the head and trunk velocities to calculate the area under the curve (AUC): The absolute values were used to compute the velocity magnitude in the balance response (aim 1) and the true values were used to compute the dominant velocity direction (aim 2).

To highlight the preferred direction of the upper body velocities in response to perturbations, we separated the ML and AP velocity components (aim 2). We also described the direction of the upper body velocity in relation to the direction of the induced loss of balance (i.e., perturbation direction), such that velocities were increased in the direction of the loss of balance (positive values) or in the opposite direction to the loss of balance (negative values). For example, right surface translation induces a leftward loss of balance. If upper body velocity in response to perturbation was towards the left, it was characterized as positive velocity in the direction of the induced loss of balance.

Perturbation direction was defined based on whether the surface translation was towards the participant’s dominant or non-dominant side. Finally, the perturbations were grouped into deciles according to the percentile of gait cycle in which they were introduced to control for different perturbation timing.

#### Statistical analysis

All statistical analyses were performed using the ‘lme4’ package in R version 4.2.2. Dependent variables comprised: MoS_AP, MoS_ML, and the AUC of head and trunk velocities. Normality of variable distributions was assessed using distribution plots and the Shapiro-Wilk test on model residuals. Outliers were examined with the ‘check_outliers’ function using "Z-scores," "ci," and "iqr" methods, identifying possible outliers for each outcome variable (see Table S2 in the supplementary materials). As outliers did not affect the models, all samples were retained, and potential outliers were not removed. For aim 1, examining the effect of the ‘arm restriction condition’ on dynamic stability and upper body velocities, Z-scores were standardized to ensure scaled coefficients across models. Then, we calculated four mixed-effect models, each representing one outcome variable, with ‘participants’ and ‘decile of gait cycle in which the perturbation was introduced’ as the *random* effects. Within-subject variables included: ‘arm restriction condition’, and ‘perturbation direction’ (surface translation toward the dominant vs. non-dominant side). Post-hoc analysis was conducted with Holm’s (sequential Bonferroni) method correction for multiple comparisons.

For aim 2, characterizing upper body kinematics in response to lateral surface translations under different arm restriction conditions, two mixed-effect models (for the head and trunk) were used, with ’participants’ and ‘decile of gait cycle in which the perturbation was introduced’ as *random* effects. Within-subject variables included: ‘plane’ (AP vs. ML), ‘arm restriction condition’ and ‘upper body strategies’ (i.e., velocity magnitude in the direction or opposite direction of the loss of balance induced by the perturbation). Post-hoc analysis with Holm’s (sequential Bonferroni) method correction for multiple comparisons was carried out.

## Results

### Participants

Participants’ characteristics are summarized in Table 1. The dependent variables of the full dataset, arranged by participant, are summarized in Table S3, in the *supplementary material*. The initial dataset included 168 trials. Thirty trials had missing marker data that excluded them from the analyses and eight trials in which the step after the perturbation was shorter than 10 kinematic data points were discarded (see Table S4 in *supplementary materials* for summary of the discarded trials). Therefore, the final analysis included 130 observations of which 48 were in the ‘free arms’, 40 in the ‘1-arm restricted’ and 42 in the ‘2-arm restricted’ conditions.

**Table 1.**
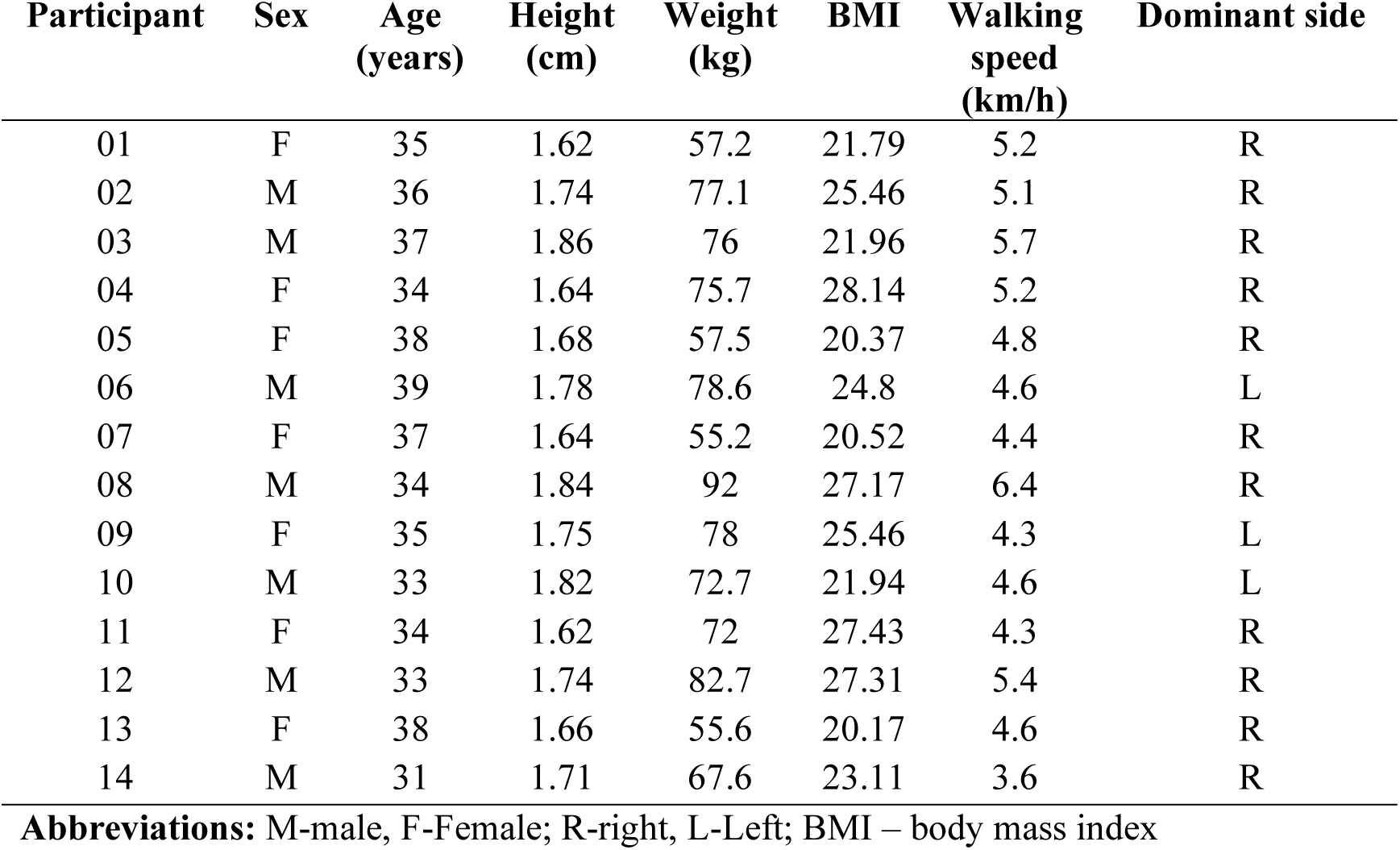
Participants’ characteristics.

| Participant | Sex | Age<br>(years) | Height<br>(cm) | Weight<br>(kg) | BMI | Walking<br>speed<br>(km/h) | Dominant side |
| --- | --- | --- | --- | --- | --- | --- | --- |
| 01 | F | 35 | 1.62 | 57.2 | 21.79 | 5.2 | R |
| 02 | M | 36 | 1.74 | 77.1 | 25.46 | 5.1 | R |
| 03 | M | 37 | 1.86 | 76 | 21.96 | 5.7 | R |
| 04 | F | 34 | 1.64 | 75.7 | 28.14 | 5.2 | R |
| 05 | F | 38 | 1.68 | 57.5 | 20.37 | 4.8 | R |
| 06 | M | 39 | 1.78 | 78.6 | 24.8 | 4.6 | L |
| 07 | F | 37 | 1.64 | 55.2 | 20.52 | 4.4 | R |
| 08 | M | 34 | 1.84 | 92 | 27.17 | 6.4 | R |
| 09 | F | 35 | 1.75 | 78 | 25.46 | 4.3 | L |
| 10 | M | 33 | 1.82 | 72.7 | 21.94 | 4.6 | L |
| 11 | F | 34 | 1.62 | 72 | 27.43 | 4.3 | R |
| 12 | M | 33 | 1.74 | 82.7 | 27.31 | 5.4 | R |
| 13 | F | 38 | 1.66 | 55.6 | 20.17 | 4.6 | R |
| 14 | M | 31 | 1.71 | 67.6 | 23.11 | 3.6 | R |

**Abbreviations:** M-male, F-Female; R-right, L-Left; BMI – body mass index

### Effect of arm restriction on gait stability and upper body velocities

Significant main effects of the ‘arm restriction condition’ were observed for MoS_ML (F_[126,2]_=4.00, p=0.021), MoS_AP (F_[126,2]_=5.97, p=0.003) and trunk velocity (F_[126,2]_=3.95, p=0.022). Notably, the ’free arms’ condition exhibited significantly lower MoS_ML compared to the ‘1-arm restricted’ condition (OR=0.40, t=2.26, p=0.026, Figure 2A); the ’free arms’ condition exhibited significantly higher MoS_AP compared to the ‘2-arm restricted’ condition (OR=-0.58, t=-3.44, p=0.001, Figure 2B); conversely, trunk velocity was significantly higher for ’2-arms restricted’ condition compared to ‘free arms’ condition (OR=0.46, t=2.75, p=0.007, Figure 2D). No significant differences between arm restriction conditions were found for head velocity (Figure 2C).

**Figure 2:**
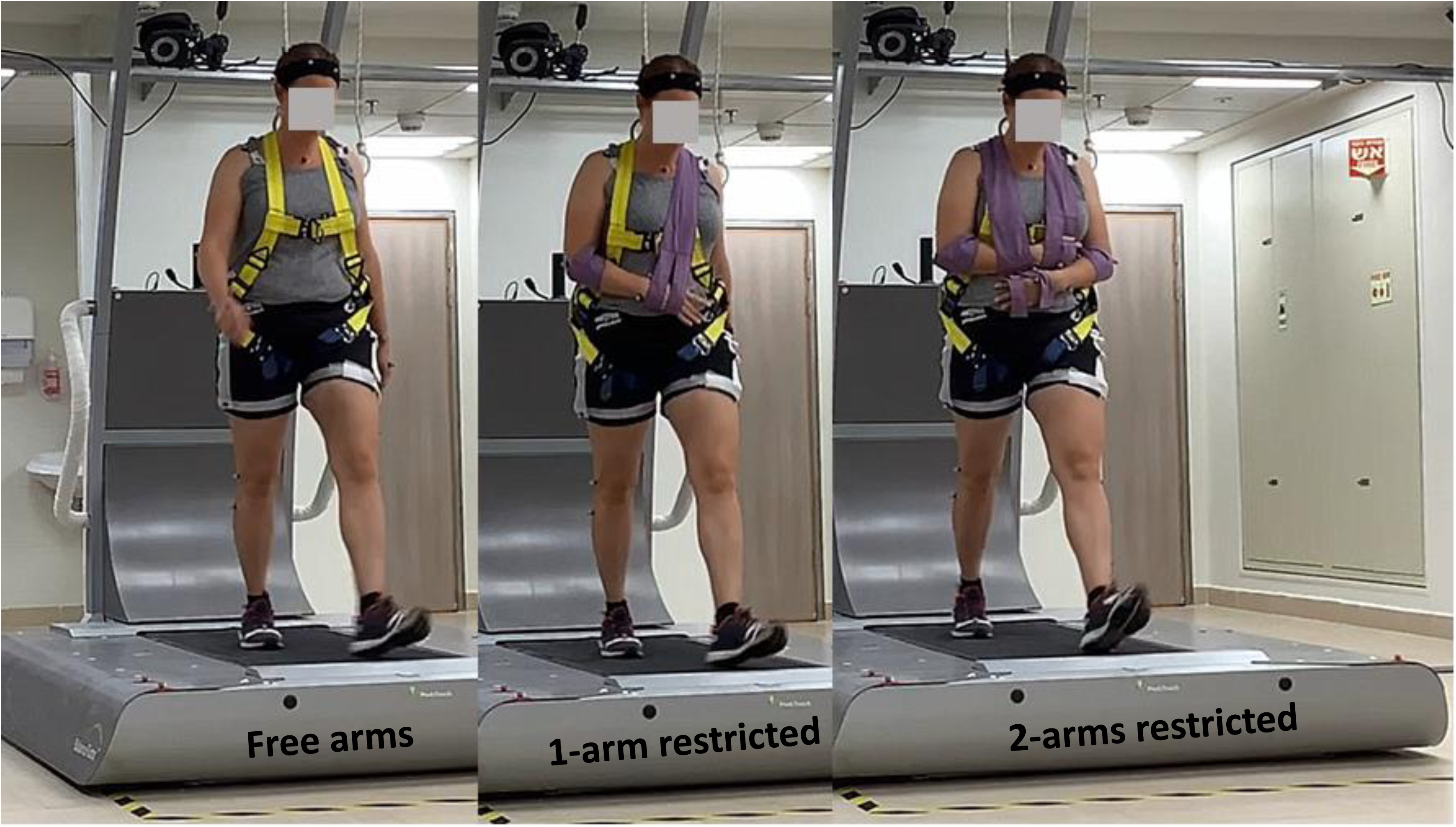
Boxplots of the effect of arm restriction condition on gait stability, i.e., margin of stability in mediolateral plane (MoS_ML, A) and in anterior-posterior plane (MoS_AP, B), head velocity (C), and trunk velocity (D). Black small dots represent actual data while larger gray dots represent the mean predicted value by the model for each variable. The black solid horizontal line represents the median, upper whiskers represent the 3rd quartile + 1.5 * Inter Quartile Range and lower whiskers represents the 1^st^ quartile - 1.5 * Inter Quartile Range. *p<0.05, **p<0.01, ***p<0.001.

### Upper body strategies

The movement of the head and trunk in the mediolateral plane was either ‘coupled’ (i.e., the velocity of both the head and trunk was in the same direction, 36.15% of responses, Figure 3A-B) or ‘decoupled’ (i.e., the velocity of the head and trunk was in opposite directions, 63.85% of responses, Figure 3C-D). In the ‘coupled’ strategy, the upper body velocities were either in the same direction as the direction of loss of balance (16.15% of responses, Figure 3A) or in the opposite direction to the loss of balance (20% of responses, Figure 3B). For the ‘decoupled’ strategy, in 37.69% of the responses the head’s velocity was in the opposite direction to the loss of balance while the trunk velocity was in the same direction as the loss of balance (Figure 3C). In 26.15% of the responses the velocity of the trunk was in the opposite direction to the loss of balance while the velocity of the head was in the same direction as the loss of balance (Figure 3D).

**Figure 3:**
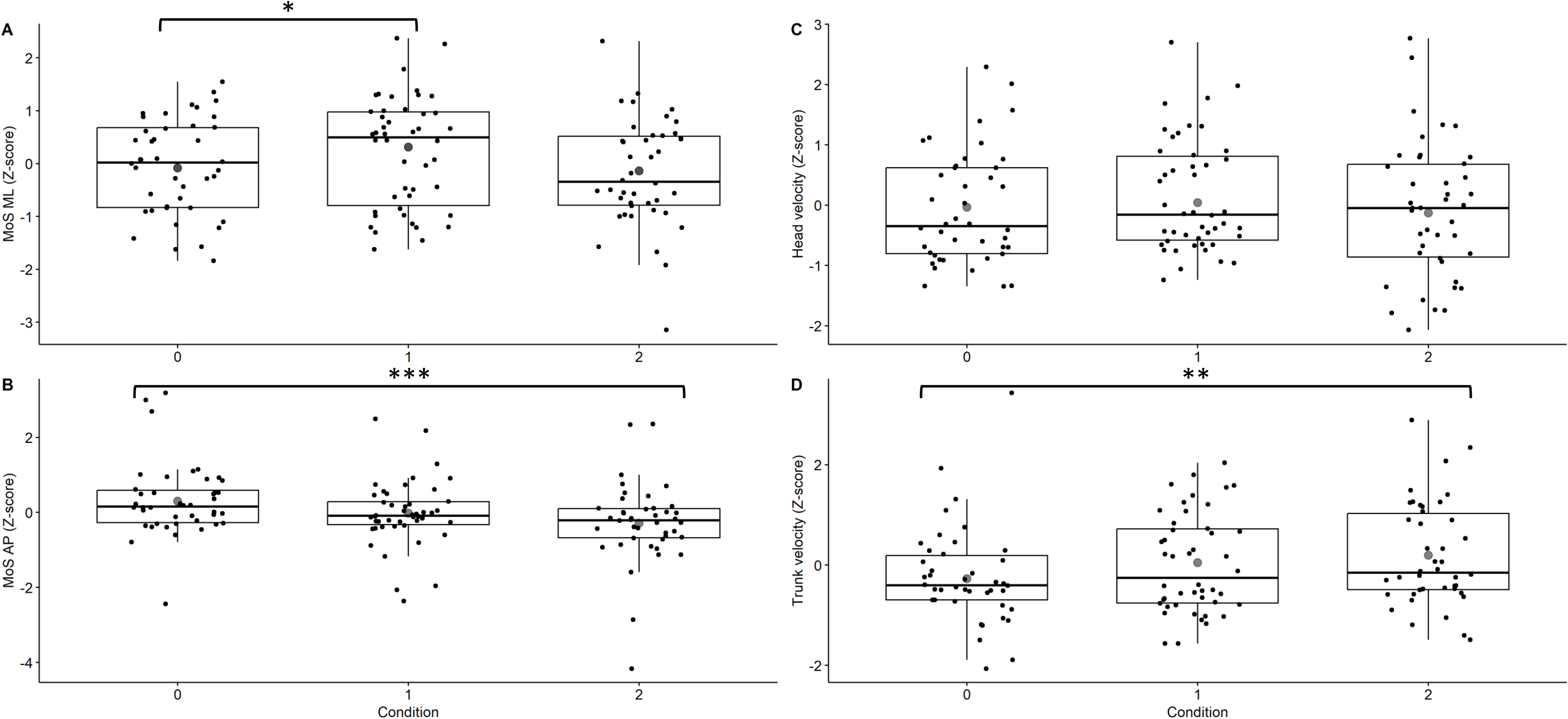
Upper body strategies in response to lateral surface translations during walking.

### Upper body kinematics

The models for the head, and trunk showed significant main effects for ‘plane’ (AP vs. ML) (F_[1,255]_=105.02, p<0.001; and F_[1,255]_=21.91, p<0.001, respectively). Head and trunk had significantly greater velocities in the ML compared to AP plane (OR=0.83, t=8.89, p<0.001 and OR=0.48, t=4.29, p<0.001, respectively, Figure 4A & 4C). We found significant main effect of ‘upper body strategies’ for the trunk (F_[1,255]_=27.72, p<0.001), indicating increased velocities in the opposite direction of the loss of balance (OR=-0.58, t=-4.64, p<0.001, Figure 4D). We didn’t find any other significant main effects (p>0.06).

**Figure 4:**
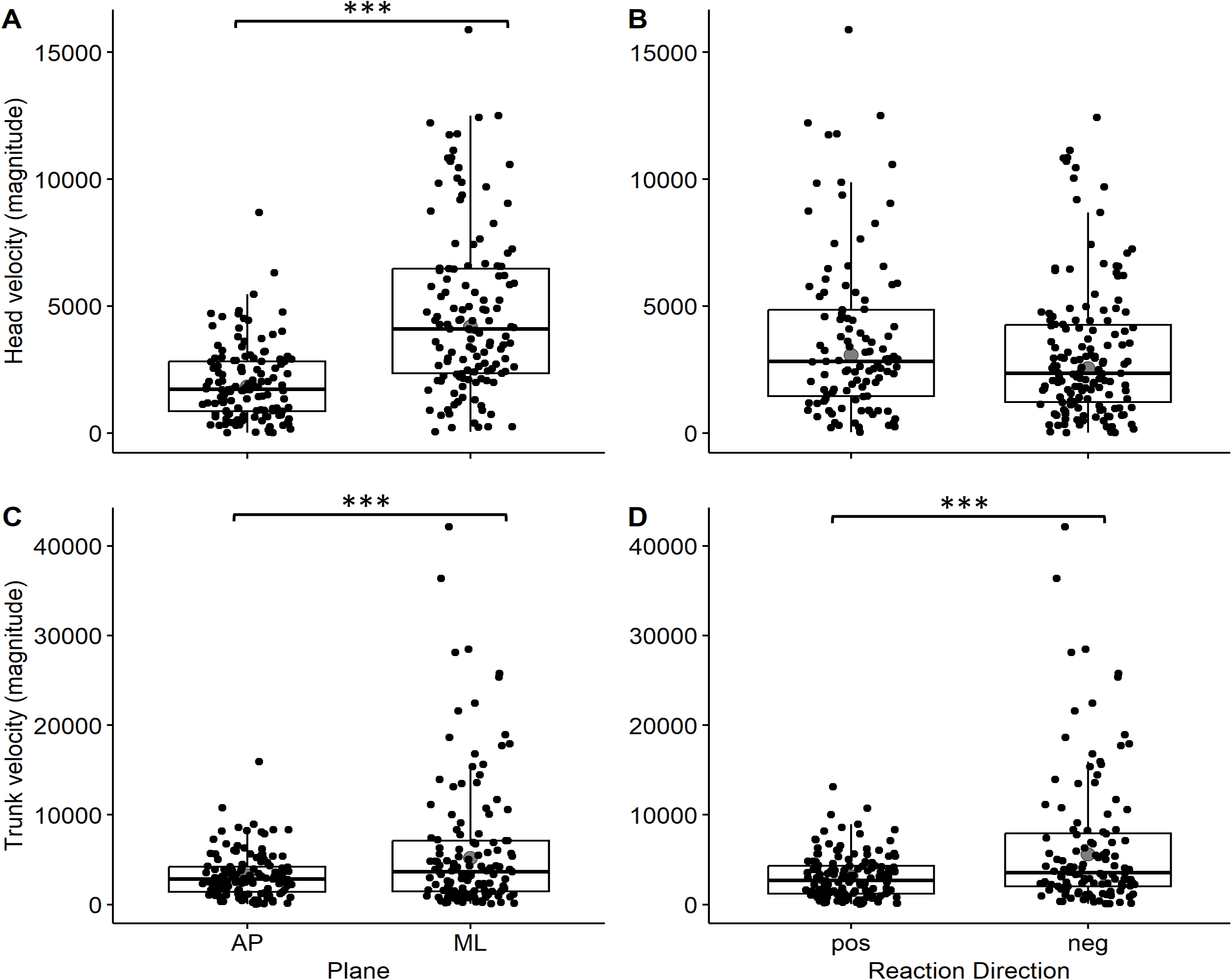
Boxplots of comparisons between upper body velocities of the head (A & B) and trunk (C & D) in the anterior-posterior (AP) vs. mediolateral (ML) planes and with (pos) or against (neg) the perturbation induced body velocity direction. black small dots represent actual data while larger gray dots represent the mean predicted value by the model for each variable. The black solid horizontal line represents the median, upper whiskers represent the 3rd quartile + 1.5 * Inter Quartile Range and lower whiskers represent the 1^st^ quartile - 1.5 * Inter Quartile Range.*P<0.05, **p<0.01, ***p<0.001.

## Discussion

In this study, we aimed to examine the effect of different arm restriction conditions on dynamic stability and upper body velocities in response to lateral surface translations. We found that 1) restricting the movements of two arms led to increased trunk velocity in comparison to ‘free arms’ condition, partially supporting our hypothesis that greater arm restriction will lead to increased upper body velocities; 2) restricting arm movements led to increased trunk velocity (but not head velocity) compared to the ‘free arms’ condition, partially supporting our hypothesis that greater arm restriction would result in increased upper body velocities; and 3) Increased head and trunk velocities were observed in the ML direction compared to the AP direction. Additionally, while the increased trunk velocities in the ML direction were opposite to the direction of the loss of balance, the increased head velocities in the ML direction were either in the same or opposite direction of the loss of balance. This partially supports our hypothesis that upper body velocities would be greater in the ML direction compared to the AP direction to counteract the loss of balance.

### Effect of arm restriction on dynamic stability and upper body velocities

Our results regarding the effect of arm restriction on dynamic stability and upper body velocities in response to lateral surface translations during walking, are consistent with previous literature. These show a significant effect of arm restriction on dynamic stability in response to balance perturbations during walking (14,41) and standing (42). A previous study showed that higher incidence of falls in response to a slip were observed in ‘arms restricted’ condition compared with ‘free arms’ condition (41). In another study reduced gait stability was demonstrated in a condition where the arm swing was restricted (14). It is of importance to note that in these studies both arms were restricted. These observations were supported by our findings of significant effect of arm restriction on dynamic stability. Our finding that the effect of arm restriction on trunk velocity was significant only in the ‘2-arm restricted’ condition but not in the ‘1-arm restricted’ condition; and that ’free arms’ condition exhibited significantly lower MoS_ML compared to the ‘1-arm restricted’ condition but not in the 2-arms condition suggests that young healthy adults can successfully compensate for not being able to react with one arm by increasing their MoS. However, in case of a substantial arm restriction (i.e., 2 arms), even young healthy adults need to change their response and react by increasing trunk velocity. This raises the question whether older adults or people with neurological disorders will demonstrate a lower “restriction threshold” to evoke a compensatory response of increased head and trunk velocities. Further research is needed to answer these questions. Furthermore, the trend of increased velocities with increased difficulty level suggests that arm restriction could be used as a methodology to focus perturbation-based balance training on trunk responses during walking.

### Upper body kinematics and strategies

Our results showed that the head and trunk velocities were significantly greater in the ML direction compared to the AP direction when controlling for the ‘arm restriction condition’ and ‘upper body strategy’. Additionally, trunk velocity was associated with upper body strategy demonstrating higher velocity in the direction opposite to the direction of loss of balance compared to the same direction as loss of balance.

The findings of two distinct upper body strategies in response to lateral perturbations, namely in the direction of the perturbation induced loss of balance direction or in the opposite direction to it are in line with previous literature (19). While previous studies demonstrated arm velocities in the direction of the loss of balance in older adults, we show for the first time this behavior in young adults in the context of head and trunk kinematics. We assume that in young adults this behavior of responding with the head and trunk in the same direction as the loss of balance without falling is made possible by compensation of other balance response mechanisms such as arm movements and stepping responses. Nevertheless, testing this assumption requires further study. Finally, our results suggest that the head and trunk, many times referred to as HAT in the literature, are most dominantly affected by the trunk movement. Moreover, the data suggests that the head and trunk should be studied separately in the context of recovery from perturbations during walking, since apparently, on some occasions, their velocities are in different directions, alluding to different calculations by the central nervous system eliciting separate motor programs.

### Limitations

Several methodological constraints should be acknowledged. First, the timing of perturbations relative to the gait cycle phase could not have been controlled for in the paradigm due to technical constraints, thus perturbations were introduced in different phases during the gait cycle. To control for that we included the timing of the perturbations defined by it disentitle of the gait cycle as a random effect variable in our models. Also, we made sure that we had at least 10 data points after the perturbation which assured lower variation of the perturbation onset in relation to the gait cycle.

Second, the arm restriction conditions were introduced in the same order between participants, with no randomization which potentially caused fatigue in the restriction conditions. One would expect to find reduced upper body velocities if fatigue was introduced, however, we did not observe that in the results. On the other hand, one may argue that fatigue may affect lower limb responses, leading to increased upper body velocities for compensation. However, in this case we should have observed negative correlations between parameters of step length and width and the upper body velocities (i.e., reduced MoS would associate with increased upper body velocities) but this was not demonstrated (see Figure S1 in *supplementary materials*). The lack of condition randomization could also have affected our results showing the ’free arms’ condition exhibited significantly lower MoS_ML compared to the ‘1-arm restricted’ condition but not in the 2-arms condition, which was introduced as the last condition. This might be a result of the participants getting used to having their hands restricted and learning of the task. Future research should ensure randomization of conditions to rule out this possibility. Finally, our study was based on healthy young adults and thus our findings cannot be generalized to other populations such as older adults and people with movement disorders. Learning about the compensatory strategies in these populations requires a separate investigation.

### Conclusions

Our findings highlight the pivotal role of increased upper body velocities as a compensatory mechanism for maintaining balance control in response to lateral surface translations during walking with arm restriction. In the current study, we observed that young adults use their trunk movement as a primary mechanism to regain balance from lateral perturbations during walking. Importantly, our observations challenge the assumption that in young adults, upper body velocities invariably oppose the direction of balance loss (see Figure 4). Additionally, the coupled velocities of the head and trunk in the direction of balance loss suggest that while increased upper body velocity is employed to maintain balance, other mechanisms, such as widening the base of support, may also be involved. The interplay between these different mechanisms requires further research. Finally, our findings challenge the traditional approach of studying the head and trunk as a single unit (HAT). Our results suggest that these are two entities that are sometimes treated differently by the motor system, leading to dominant velocities in opposite directions.

## Ethics approval and consent to participate

This study was approved by the Helsinki committee at Loewenstein medical rehabilitation center, Israel (Approval Number 0021-14-LOE). All participants gave signed, written, informed consent.

## Availability of data materials

The raw datasets used and/or analyzed during the current study are available from the corresponding author on request. A table summarizing individualized mean gait spatiotemporal parameters and a table summarizing the median values and inter quartile range of individualized preprocessed data used for the analysis are provided in Table S1 and Table S3 in the Supplementary materials.

## Author’s contribution

Conceived and designed the experiments: AH, SH. Conducted the experiments and data collection: AL, AF, AH, SH. Analyzed the results: UR, AL, AH, YP. Participated in data interpretation and writing the final version of the manuscript: UR, AL, AF, YP, AH, SH. All authors have reviewed and edited the manuscript and have read and approved the final manuscript.

## References

1. Wang S, Bhatt T. Kinematic Measures for Recovery Strategy Identification following an Obstacle-Induced Trip in Gait. J Mot Behav. 2023 Jan 5;55(2):193–201.

2. Wang S, Pai Y-C, Bhatt T. Neuromuscular mechanisms of motor adaptation to repeated gait-slip perturbations in older adults. Sci Rep. 2022 Nov 18;12(1):19851.

3. Huntley AH, Rajachandrakumar R, Schinkel-Ivy A, Mansfield A. Characterizing slip-like responses during gait using an entire support surface perturbation: Comparisons to previously established slip methods. Gait Posture. 2019 Mar;69:130–5.

4. Rosenblum U, Melzer I, Zeilig G, Plotnik M. Muscle activation profile is modulated by unexpected balance loss in walking. Gait Posture. 2022 Mar;93:64–72.

5. Batcir S, Shani G, Shapiro A, Melzer I. Characteristics of step responses following varying magnitudes of unexpected lateral perturbations during standing among older people - a cross-sectional laboratory-based study. BMC Geriatr. 2022 May 6;22(1):400.

6. MacKinnon CD, Winter DA. Control of whole body balance in the frontal plane during human walking. J Biomech. 1993 Jun;26(6):633–44.

7. Hirasaki E, Kubo T, Nozawa S, Matano S, Matsunaga T. Analysis of head and body movements of elderly people during locomotion. Acta Otolaryngol Suppl. 1993;501:25–30.

8. Pozzo T, Berthoz A, Lefort L. Head stabilization during various locomotor tasks in humans. I. Normal subjects. Exp Brain Res. 1990;82(1):97–106.

9. Pozzo T, Levik Y, Berthoz A. Head and trunk movements in the frontal plane during complex dynamic equilibrium tasks in humans. Exp Brain Res. 1995;106(2):327–38.

10. Nakakubo S, Doi T, Sawa R, Misu S, Tsutsumimoto K, Ono R. Does arm swing emphasized deliberately increase the trunk stability during walking in the elderly adults? Gait Posture. 2014 Sep;40(4):516–20.

11. Punt M, Bruijn SM, Wittink H, van Dieën JH. Effect of arm swing strategy on local dynamic stability of human gait. Gait Posture. 2015 Feb;41(2):504–9.

12. Angelini L, Damm P, Zander T, Arshad R, Di Puccio F, Schmidt H. Effect of arm swinging on lumbar spine and hip joint forces. J Biomech. 2018 Mar 21;70:185–95.

13. Tokur D, Grimmer M, Seyfarth A. Review of balance recovery in response to external perturbations during daily activities. Hum Mov Sci. 2020 Feb;69:102546.

14. Bruijn SM, Meijer OG, Beek PJ, van Dieën JH. The effects of arm swing on human gait stability. J Exp Biol. 2010 Dec 1;213(Pt 23):3945–52.

15. Pijnappels M, Kingma I, Wezenberg D, Reurink G, van Dieën JH. Armed against falls: the contribution of arm movements to balance recovery after tripping. Exp Brain Res. 2010 Apr;201(4):689–99.

16. Gholizadeh H, Hill A, Nantel J. Effect of arm motion on postural stability when recovering from a slip perturbation. J Biomech. 2019 Oct 11;95:109269.

17. Maki BE, McIlroy WE. The role of limb movements in maintaining upright stance: the “change-in-support” strategy. Phys Ther. 1997 May;77(5):488– 507.

18. Marigold DS, Bethune AJ, Patla AE. Role of the unperturbed limb and arms in the reactive recovery response to an unexpected slip during locomotion. J Neurophysiol. 2003 Apr;89(4):1727–37.

19. Alissa N, Akinlosotu RY, Shipper AG, Wheeler LA, Westlake KP. A systematic review of upper extremity responses during reactive balance perturbations in aging. Gait Posture. 2020 Oct;82:138–46.

20. Roos PE, McGuigan MP, Kerwin DG, Trewartha G. The role of arm movement in early trip recovery in younger and older adults. Gait Posture. 2008 Feb;27(2):352–6.

21. Allum JHJ, Carpenter MG, Honegger F, Adkin AL, Bloem BR. Age-dependent variations in the directional sensitivity of balance corrections and compensatory arm movements in man. J Physiol (Lond). 2002 Jul 15;542(Pt 2):643–63.

22. Troy KL, Donovan SJ, Grabiner MD. Theoretical contribution of the upper extremities to reducing trunk extension following a laboratory-induced slip. J Biomech. 2009 Jun 19;42(9):1339–44.

23. Lee-Confer JS, Bradley NS, Powers CM. Quantification of reactive arm responses to a slip perturbation. J Biomech. 2022 Mar;133:110967.

24. Merrill Z, Chambers AJ, Cham R. Arm reactions in response to an unexpected slip-Impact of aging. J Biomech. 2017 Jun 14;58:21–6.

25. Bruijn SM, Sloot LH, Kingma I, Pijnappels M. Contribution of arm movements to balance recovery after tripping in older adults. J Biomech. 2022 Mar;133:110981.

26. Maki BE, Edmondstone MA, McIlroy WE. Age-related differences in laterally directed compensatory stepping behavior. J Gerontol A Biol Sci Med Sci. 2000 May;55(5):M270–7.

27. Mille M-L, Johnson ME, Martinez KM, Rogers MW. Age-dependent differences in lateral balance recovery through protective stepping. Clin Biomech (Bristol, Avon). 2005 Jul;20(6):607–16.

28. Hilliard MJ, Martinez KM, Janssen I, Edwards B, Mille M-L, Zhang Y, et al. Lateral balance factors predict future falls in community-living older adults. Arch Phys Med Rehabil. 2008 Sep;89(9):1708–13.

29. Hof AL, Gazendam MGJ, Sinke WE. The condition for dynamic stability. J Biomech. 2005 Jan;38(1):1–8.

30. Bruijn SM, Meijer OG, Beek PJ, van Dieën JH. Assessing the stability of human locomotion: a review of current measures. J R Soc Interface. 2013 Jun 6;10(83):20120999.

31. Siragy T, Nantel J. Quantifying dynamic balance in young, elderly and parkinson’s individuals: A systematic review. Front Aging Neurosci. 2018 Nov 22;10:387.

32. König M, Epro G, Seeley J, Potthast W, Karamanidis K. Retention and generalizability of balance recovery response adaptations from trip perturbations across the adult life span. J Neurophysiol. 2019 Nov 1;122(5):1884–93.

33. Kagawa T, Suzuki R. Balance map analysis for visualization and quantification of balance in human walking. IEEE Trans Neural Syst Rehabil Eng. 2021 Oct 28;29:2153–63.

34. Gerards MHG, Meijer K, Karamanidis K, Grevendonk L, Hoeks J, Lenssen AF, et al. Adaptability to balance perturbations during walking as a potential marker of falls history in older adults. Front Sports Act Living. 2021 May 19;3:682861.

35. Winter DA. Human balance and posture control during standing and walking. Gait Posture. 1995 Dec;3(4):193–214.

36. Kuo AD. Stabilization of Lateral Motion in Passive Dynamic Walking. Int J Rob Res. 1999 Sep 1;18(9):917–30.

37. Townsend MA. Biped gait stabilization via foot placement. J Biomech. 1985;18(1):21–38.

38. Zeni JA, Richards JG, Higginson JS. Two simple methods for determining gait events during treadmill and overground walking using kinematic data. Gait Posture. 2008 May;27(4):710–4.

39. Madehkhaksar F, Klenk J, Sczuka K, Gordt K, Melzer I, Schwenk M. The effects of unexpected mechanical perturbations during treadmill walking on spatiotemporal gait parameters, and the dynamic stability measures by which to quantify postural response. PLoS ONE. 2018 Apr 19;13(4):e0195902.

40. McIlroy WE, Maki BE. Early activation of arm muscles follows external perturbation of upright stance. Neurosci Lett. 1995 Jan 30;184(3):177–80.

41. Lee-Confer JS, Kulig K, Powers CM. Constraining the arms during a slip perturbation results in a higher fall frequency in young adults. Hum Mov Sci. 2022 Dec;86:103016.

42. Cheng KB, Huang Y-C, Kuo S-Y. Effect of arm swing on single-step balance recovery. Hum Mov Sci. 2014 Dec;38:173–84.

