## Supplementary Materials for "The Effect of Arm Restriction on Dynamic Stability and Upper Body Responses to Lateral Loss of Balance During Walking: An Observational Study"

### Methods

**Table S1:** Summary of Gait spatiotemporal parameters by participant and condition

| id | condition | Step Velocity (mean) | Step Length (mean) | Step Width (mean) | Step Time (mean) |
|---|---|---|---|---|---|
| 1 | free arms | 1.035064 | 0.498677 | 0.129329 | 0.475 |
| 1 | 1-arm restricted | 1.014932 | 0.459138 | 0.153165 | 0.445833 |
| 1 | 2-arms restricted | 1.293031 | 0.611123 | 0.063122 | 0.470833 |
| 2 | free arms | 0.601664 | 0.28984 | 0.210631 | 0.445833 |
| 2 | 2-arms restricted | 0.52922 | 0.211599 | 0.256685 | 0.3875 |
| 3 | free arms | 1.025241 | 0.494495 | 0.151868 | 0.466667 |
| 3 | 1-arm restricted | 1.217329 | 0.609416 | 0.070635 | 0.5 |
| 3 | 2-arms restricted | 0.988187 | 0.474084 | 0.08013 | 0.470833 |
| 4 | free arms | 0.528792 | 0.319603 | 0.184809 | 0.920833 |
| 4 | 1-arm restricted | 1.062115 | 0.436534 | 0.060547 | 0.410417 |
| 4 | 2-arms restricted | 1.04799 | 0.443166 | 0.159795 | 0.416667 |
| 5 | free arms | 0.660191 | 0.318776 | 0.262913 | 0.483333 |
| 5 | 1-arm restricted | 0.776788 | 0.401557 | 0.162622 | 0.510417 |
| 5 | 2-arms restricted | 1.270402 | 0.591453 | 0.070472 | 0.466667 |
| 6 | free arms | 0.372136 | 0.152975 | 0.251207 | 0.40625 |
| 6 | 1-arm restricted | 0.918072 | 0.370888 | 0.22595 | 0.404167 |
| 6 | 2-arms restricted | 0.580313 | 0.4739 | 0.166724 | 0.814583 |
| 7 | free arms | 0.663487 | 0.390376 | 0.188742 | 0.839583 |
| 7 | 1-arm restricted | 1.179192 | 0.490203 | 0.249039 | 0.427083 |
| 8 | free arms | 1.297004 | 0.714315 | 0.0835 | 0.639583 |
| 8 | 1-arm restricted | 1.343247 | 0.644 | 0.081484 | 0.483333 |
| 8 | 2-arms restricted | 1.484243 | 0.667623 | 0.09948 | 0.447917 |
| 9 | free arms | 0.659943 | 0.3239 | 0.179707 | 0.470833 |
| 10 | free arms | 0.842356 | 0.429599 | 0.150922 | 0.508333 |
| 10 | 1-arm restricted | 0.669028 | 0.308586 | 0.229232 | 0.447917 |
| 10 | 2-arms restricted | 1.006051 | 0.367078 | 0.264252 | 0.385417 |
| 11 | 1-arm restricted | 0.703484 | 0.407579 | 0.264161 | 0.616667 |
| 11 | 2-arms restricted | 0.454027 | 0.184861 | 0.14891 | 0.389583 |
| 12 | free arms | 1.031209 | 0.513268 | 0.127415 | 0.491667 |
| 12 | 1-arm restricted | 0.935105 | 0.54912 | 0.131887 | 0.64375 |
| 12 | 2-arms restricted | 1.011698 | 0.450462 | 0.143607 | 0.439583 |
| 13 | free arms | 0.886571 | 0.425801 | 0.14517 | 0.475 |
| 13 | 1-arm restricted | 0.845785 | 0.449081 | 0.044498 | 0.529167 |
| 13 | 2-arms restricted | 0.904316 | 0.567004 | 0.149039 | 0.779167 |
| 14 | free arms | 0.666427 | 0.500521 | 0.192385 | 0.78125 |
| 14 | 1-arm restricted | 0.843013 | 0.427372 | 0.292243 | 0.527083 |
| 14 | 2-arms restricted | 0.901507 | 0.534707 | 0.134317 | 0.608333 |

**Table S2:** summary of possible outliers

| Case | id | condition | Perturbation #, direction | outcome measure | Value |
|---|---|---|---|---|---|
| 1 | 1 | 2-arms restricted | 1, non-dominant | MOS_AP | -2.58E-01 |
| 2 | 5 | free arms | 2, non-dominant | MOS_AP | 1.60E-01 |
| 3 | 5 | free arms | 3, non-dominant | Trunk velocity | 4.44E+04 |
| 4 | 1 | 2-arms restricted | 1, non-dominant | Dominant shoulder velocity | 1.91E+04 |
| 5 | 4 | 2-arms restricted | 1, dominant | Non-dominant shoulder velocity | 2.51E+04 |

MoS_AP = margin of stability in anterior-posterior direction.

### Results

**Table S3:** Summary of dynamic balance and upper body velocities by participant and condition

| ID | Condition | Mos_ML, median(IQR) | Mos_AP, median(IQR) | Head, median(IQR) | Trunk, median(IQR) |
|---|---|---|---|---|---|
| 1 | 2-arms restricted | 0.09(0.01) | -0.04(0.02) | 3.59E+03(1.37E+03) | 8.98E+03(1.57E+03) |
| 1 | 1-arm restricted | 0.09(0.02) | -0.03(0.01) | 3.57E+03(5.53E+02) | 9.11E+03(2.20E+03) |
| 1 | free arms | 0.11(0.01) | 0.00(0.01) | 4.07E+03(1.41E+03 | 9.44E+03(2.33E+03) |
| 2 | 2-arms restricted | 0.12(0.01) | -0.02(0.00) | 1.42E+03(2.14E+02 | 6.62E+03(1.08E+03) |
| 2 | free arms | 0.13(0.01) | 0.01(0.00) | 2.93E+03(1.51E+03) | 9.27E+03(3.00E+03) |
| 3 | 2-arms restricted | 0.13(0.01) | -0.03(0.01) | 5.56E+03(1.24E+03) | 6.46E+03(1.54E+03) |
| 3 | 1-arm restricted | 0.14(0.03) | -0.03(0.00) | 6.08E+03(1.36E+03) | 6.76E+03(2.44E+03) |
| 3 | free arms | 0.12(0.07) | -0.03(0.01) | 6.03E+03(1.45E+03) | 5.31E+03(3.28E+03) |
| 4 | 2-arms restricted | 0.07(0.05) | -0.04(0.05) | 4.49E+0(34.96E+02) | 1.07E+04(4.02E+03) |
| 4 | 1-arm restricted | 0.14(0.02) | -0.03(0.03) | 5.67E+03(9.46E+02) | 1.05E+04(1.14E+03) |
| 4 | free arms | 0.06(0.00) | -0.02(0.02) | 4.08E+03(2.51E+03) | 7.28E+03(4.36E+03) |
| 5 | 2-arms restricted | 0.10(0.01) | -0.07(0.04) | 1.01E+04(6.02E+03) | 2.58E+04(1.47E+04) |
| 5 | 1-arm restricted | 0.14(0.03) | 0.01(0.01) | 3.80E+03(1.10E+03) | 9.82E+03(3.06E+03) |
| 5 | free arms | 0.11(0.03) | -0.10(0.06) | 2.43E+03(1.95E+02) | 6.16E+03(1.09E+03) |
| 6 | 2-arms restricted | 0.12(0.05) | -0.03(0.03) | 2.24E+03(1.50E+02) | 7.89E+03(4.17E+02) |
| 6 | 1-arm restricted | 0.18(0.04) | -0.04(0.05) | 2.88E+03(1.97E+03) | 1.10E+04(1.28E+04) |
| 6 | free arms | 0.13(0.05) | -0.01(0.00) | 2.72E+03(6.39E+02) | 8.58E+03(2.18E+03) |
| 7 | 1-arm restricted | 0.14(0.03) | -0.08(0.14) | 5.14E+03(2.99E+03) | 1.37E+04(7.75E+03) |
| 7 | free arms | 0.11(0.02) | 0.03(0.02) | 4.34E+03(2.07E+03) | 1.03E+04(4.13E+03) |
| 8 | 2-arms restricted | 0.13(0.01) | -0.07(0.01) | 3.52E+03(4.49E+02) | 1.30E+04(7.50E+02) |
| 8 | 1-arm restricted | 0.14(0.01) | -0.06(0.03) | 3.96E+03(6.68E+02) | 1.35E+04(2.40E+03) |
| 8 | free arms | 0.10(0.02) | -0.05(0.01) | 3.26E+03(1.01E+03) | 1.14E+04(3.69E+03) |
| 9 | free arms | 0.11(0.03) | 0.03(0.01) | 1.20E+03(7.53E+02) | 2.31E+03(1.41E+03) |
| 10 | 2-arms restricted | 0.12(0.09) | -0.13(0.013) | 7.40E+03(1.20E+04) | 2.28E+04(3.10E+04) |
| 10 | 1-arm restricted | 0.11(0.09) | -0.03(0.02) | 2.91E+03(2.37E+03) | 7.32E+03(4.62E+03( |
| 10 | free arms | 0.15(0.03) | -0.02(0.01) | 3.15E+03(1.60E+03) | 7.63E+03(2.26E+03) |
| 11 | 2-arms restricted | 0.09(0.02) | -0.01(0.02) | 4.83E+03(1.39E+03) | 8.60E+03(7.07E+02) |
| 11 | 1-arm restricted | 0.10(0.01) | -0.01(0.01) | 3.42E+03(6.73E+02) | 5.52E+03(1.16E+03) |
| 12 | 2-arms restricted | 0.12(0.04) | -0.03(0.04) | 4.07E+03(4.33E+02) | 1.27E+04(1.73E+03) |
| 12 | 1-arm restricted | 0.13(0.08) | -0.03(0.01) | 5.62E+03(9.45E+02) | 1.06E+04(1.37E+03) |
| 12 | free arms | 0.12(0.08) | -0.04(0.00) | 4.00E+03(3.34E+03) | 9.12E+03(4.25E+03) |
| 13 | 2-arms restricted | 0.08(0.02) | -0.04(0.03) | 1.33E+04(9.78E+03) | 2.30E+04(1.71E+04) |
| 13 | 1-arm restricted | 0.11(0.06) | 0.03(0.02) | 4.09E+03(3.14E+02) | 8.91E+03(2.89E+02) |
| 13 | free arms | 0.11(0.05) | 0.03(0.01) | 1.81E+03(3.44E+03) | 3.97E+03(8.05E+03) |
| 14 | 2-arms restricted | 0.13(0.01) | 0.07(0.08) | 6.60E+03(2.22E+03) | 1.62E+04(5.67E+03) |
| 14 | 1-arm restricted | 0.14(0.02) | 0.11(0.07) | 3.02E+03(1.98E+03) | 6.47E+03(4.87E+03) |
| 14 | free arms | 0.14(0.01) | 0.14(0.05) | 5.80E+03(3.33E+03) | 1.26E+04(1.07E+04) |

MoS_AP = margin of stability in anterior-posterior direction; MoS_ML = margin of stability in mediolateral direction; IQR = inter quartile range.

**Table S4:** summary of trials where the step after the perturbation was shorter than 10 kinematic data points

| Case | id | condition | Perturbation #,direction | Number of samples |
|---|---|---|---|---|
| 1 | 4 | free arms | 2, non-dominant | 7 |
| 2 | 9 | free arms | 2, dominant | 5 |
| 3 | 10 | free arms | 1, non-dominant | 6 |
| 4 | 10 | 1-arm restricted | 1, non-dominant | 2 |
| 5 | 12 | free arms | 3, non-dominant | 4 |
| 6 | 13 | free arms | 2, non-dominant | 2 |
| 7 | 13 | free arms | 4, dominant | 1 |
| 8 | 14 | 1-arm restricted | 4, dominant | 5 |

### Discussion

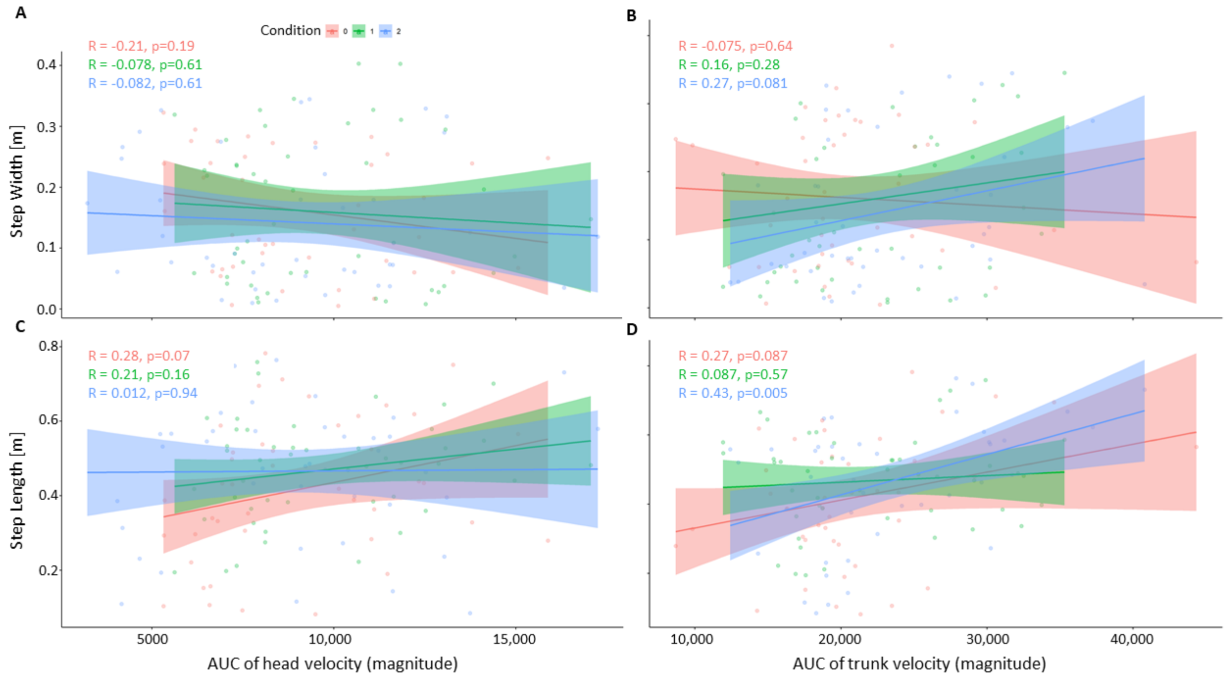


**Figure S1:** The relationship between dynamic balance and upper body velocities (A-D) in the different arm restriction conditions (red dots – 'free arms', green dots – '1-arm restricted' and blue dots – '2-arms restricted'. Solid black line represents the regression line and shaded area around it the 95% confidence interval. Abbreviations: MoS, margin of stability; ML, mediolateral; AP, anterior-posterior.
